# The effects of phenotypic plasticity on diversification rates and adaptive evolution in simulated environments with different climatic and cost contexts

**DOI:** 10.1101/2023.12.30.573714

**Authors:** Emerson Campos Barbosa Júnior, Pavel Dodonov, Hilton F. Japyassú, Bruno Vilela

**Affiliations:** Instituto de Biologia, Universidade Federal da Bahia, CEP 40170-110, Salvador, Bahia, Brazil

**Keywords:** anthropogenic changes, contemporary adaptation, extended evolutionary synthesis, macroecology, macroevolution, natural selection

## Abstract

Phenotypic plasticity can either hinder or facilitate genetic evolution, thus affecting macroevolution. However, the mechanisms and associations of phenotypic plasticity with biodiversity patterns remain unresolved. In this paper, we investigate the effect of phenotypic plasticity on adaptive evolution in the context of climatic changes and plasticity costs, specifically examining the rates of trait evolution, speciation, extinction, and diversification. We employed an eco-evolutionary agent-based model, incorporating body temperature as a plastic trait that dynamically responds to fluctuations in environmental temperature. We found the same pattern of results for both climate and plasticity cost contexts. We found that an increase in plasticity leads to a decrease in extinction and trait evolution rates, with high plasticity serving to postpone mass extinction events. Systems with plastic species experience higher speciation rates. Additionally, as plasticity increases in systems, diversification rates also increase. Our study supports the hypothesis that plasticity can facilitate adaptive evolution. It provides new insights into the effect of plasticity on adaptive evolution at larger scales, considering relevant mechanisms and factors such as population size, grouping, occupancy, plasticity extension, and dispersion.

## Introduction

In traditional evolutionary synthesis, phenotypic plasticity, which refers to the capacity of a particular genotype to generate different phenotypes in response to varying environmental conditions (Pigliucci, 2001), was not considered as a significant factor for the future adaptation of species (Nicoglou, 2014). However, this has recently changed, and there is increasing recognition of the potential of plasticity in altering the selective and adaptive processes of traits (Laland, 2015; Ghalambor et al. 2015; Wong and Candolin 2015; Fox et al. 2019; Scheiner and Mindell, 2020). Nevertheless, it is not yet clear how phenotypic plasticity affects adaptive evolution (Wong and Candolin 2015; Kelly 2019; Fox et al. 2019), with evidence indicating that it can both hamper or facilitate genetic evolution (Ghalambor et al. 2007; Wong and Candolin 2015).

Phenotypic plasticity can hinder adaptation by promoting the dominance of a particular phenotype in the population’s genetic pool (Scheiner, 2020). This can lead to a decrease in genetic variation by selecting a single genotype group, thus homogenizing the population and resulting in stabilizing selection. Under stabilizing selection, there would be continuous selection of the dominant adaptive phenotype with a similar genotype (Ghalambor et al., 2007; Wong and Candolin, 2015; Scheiner, 2020). This dominance can reduce the directionality of selection and hide the underlying genetic variation, acting as a shield that prevents other adaptive phenotypes or genotypes from being selected for that environment (Ghalambor et al., 2007; Pfennig et al., 2010; Wong and Candolin, 2015; Scheiner, 2020).

Alternatively, plasticity can facilitate adaptation by either reducing extinction and prolonging the time for adaptive evolution to occur or through canalization via the genetic assimilation of a phenotype with optimal fitness (Baldwin 1896; Ghalambor et al. 2015; Scheiner, 2020; Pfenning, 2021). The first facilitation hypothesis is called the “buying time hypothesis”, where plasticity works as a shield against environmental changes, promoting population persistence (indirect facilitation) (Baldwin 1896). The second facilitation hypothesis, known as the “Plasticity-First Hypothesis”, involves genetic accommodation and assimilation. Genetic accommodation is an evolutionary mechanism in which a novel phenotype, resulting from either a mutation or environmental perturbation, undergoes refinement into an adaptive phenotype through a series of quantitative genetic changes (Ehrenreich and Pfennig, 2016). This process can lead to a plasticity decrease known as “genetic assimilation,” whereby a phenotypic character becomes taken over by the genotype, and sometimes there is a complete plasticity loss referred to as “canalization” (Waddington, 1942; Ehrenreich and Pfennig, 2016).

The “Plasticity-First Hypothesis” suggests that selection operates on the phenotype in the initial step. When the environment changes, there may be an adaptive adjustment in the phenotype known as phenotypic accommodation (West-Eberhard 2003). After that, the trait would be genetically accommodated, and in some cases, canalized by genetic assimilation. This process of phenotypic and genetic accommodation would allow individuals to accumulate genetic variation and expose unexpressed genes, thereby accelerating adaptive evolution through directional selection (ecological time and direct facilitation) (West-Eberhard 2003; Ghalambor et al. 2007, 2010; Wong and Candolin 2015; Pfenning, 2021).

One of the main reasons for this is the lack of understanding of the underlying mechanisms related to the influence of plasticity on evolution and its association with current and future biodiversity patterns (Wong and Candolin, 2015; Fox et al., 2019). The difficulty in understanding the effect of plasticity on adaptive evolution is related to four main topics: (i) how trait evolution occurs and the associated mechanisms (Wong and Candolin, 2015; Bro-Jorgensen et al., 2019); (ii) how adaptive evolution unfolds in the context of macroevolution and eco-evolutionary dynamics (Bro-Jorgensen et al., 2019; Fox et al., 2019); (iii) how extinctions occur during this process (Wong and Candolin, 2015; Bro-Jorgensen et al., 2019; Fox et al., 2019); and (iv) how diversification occurs in face of human-induced rapid environmental changes (Bro-Jorgensen et al. 2019; Fox et al. 2019; Kelly 2019).

Here, we used an agent-based model to investigate the effect of phenotypic plasticity on adaptive evolution in the context of climatic changes and considering plasticity costs. Specifically, we examine the rates of trait evolution, speciation, extinction, and diversification. Our hypothesis is that phenotypic plasticity facilitates adaptive evolution by increasing the rates of trait evolution and speciation, while reducing extinction rates in all contexts. To achieve this, we incorporated plasticity in individuals’ body temperature into a spatially explicit eco-evolutionary computational simulation implemented by Hagen et al. (2021). Throughout the simulation, processes such as speciation, dispersion, extinction, and evolution unfolded over the course of a thousand steps to allow macroevolutionary changes to happen.

## Material and methods

### Model characterization

We used the agent-based modeling (ABM) framework “gen3sis”, which is implemented as an R package (R Development Core Team 2019; Hagen et al., 2021) focused on providing background code for running simulations in macroecology and macroevolution. “gen3sis” works with two inputs: landscape and configuration. Landscape contains spatiotemporal information about the environment where all processes occur, such as the world format, environmental variables/conditions (land or ocean, temperature, aridity, among others), and distance matrices (connectivity). Configuration contains general information on the simulation, including the number of species and individuals, beginning and end time of the simulation, speciation, dispersion, ecology, evolution, and others. Simulations occur in three steps: (1) preparing the landscape from a raster file and defining the configuration, (2) creating the initial world by loading information such as a random seed, the initial landscape, species distribution, and phylogenetic statistics (e.g. living and total species), using the landscape and configuration as foundation, and (3) updating the information on the world at each time step. All the code used in the simulation is available on Github (link: https://github.com/emersonjr25/PlasticityOnEvolution).

### Modeling

Our world was based on the current center of the world, with a spatial extent of latitude (-40°;40°) and longitude (-80°;80°). We used the World Center map, which is one of the default maps included in the “`gen3sis`” package (Hagen et al., 2021). The simulations were run for 1000 timesteps but were stopped once the world reached 0 or 50,000 species. The start time was 1000 timesteps and the end time was 1 (from past to current time). There were three environmental variables: temperature, aridity, and area, with temperature being the variable that changed over the 1000 timesteps. In our configuration, individuals had two traits: body temperature and dispersal. The initial abundance was one individual of one species, and the initial individual temperature values were equal to the environmental temperature. The individual can disperse, reproduce by bipartition, face death in situations of low fitness, possess identity, body temperature value, and geographic position. In the second time step, the environmental temperature began to change based on the climatic context, either through slow or fast changes. In the slow climatic context, temperature values took 1000 timesteps to transition from one extreme temperature (colder) to the other extreme (warmer). On the other hand, in fast climate changes, the temperature changed twice as fast as in the slow context, taking half the time to transition from one extreme value to another. We also introduced spatial temperature variations at all timesteps, which were based on latitudinal and longitudinal variations observed in the real world, with warmer temperatures in the center and the coldest temperatures farther away (detailed in the Supplementary Material).

The simulation consists of four core algorithms: speciation, dispersal, evolution, and ecology, which occur sequentially in each time step. Speciation is based on a divergence threshold of 10. The divergence value accumulates each time step when species become geographically distant. If the species exceed the limit, speciation occurs. To calculate the amount of speciation in each time step, we subtracted the total number of species in the step t-1 from the number of all species in the step t. Dispersal is based on drawing a value using fixed definitions that depend on the number of free cells in the world, species abundance, and geographic distance. We set a maximum of 270 free cells to dispersion in the world, which creates pressure for species to survive through adaptation or plasticity. Evolution returns traits with modified values based on a fixed evolutionary power of 0.001 for mutation. Ecology is responsible for returning abundance, which is defined by the correspondence between the trait value and the environmental temperature value. The optimal trait value is close to the environmental temperature value. Individuals die based on the distance between their internal and external temperature, representing a selective process. If all individuals of a species die, it becomes extinct. The farther away the temperature trait is from the environmental temperature, the smaller is the fitness – representing ectotherms animals (plasticity increases the range of the trait value of the species). We calculated the amount of extinction in each time step as the number of living species at step t minus the number of living species at step t-1 (excluding those that speciated from t-1 to t).

We considered body temperature as a plastic trait that varies in response to changes in environmental temperature. Plasticity in this context represents the variation in trait values around the original trait value, with the direction of body temperature variation towards the optimal trait value. To test the effect of plasticity on evolutionary rates, we used eight deterministic different ranges: 0, 0.05, 0.1, 0.15, 0.25, 0.5, 0.75, and 1. Greater plasticity was represented by a wider range of trait variation, with range values representing an increase in percentage (e.g., 0.1 = 10% increase and 1 = 100% increase). We conducted simulations in both the absence and presence of the plasticity cost. The cost of plasticity involves changing the body temperature value further away from the optimal body temperature (calculated proportionally based on the plasticity level). To determine the specific value of the plasticity cost in the presence of it, we identified the maximum value of the plasticity cost (10%) that the system could sustain without experiencing mass extinctions within the initial timesteps.

We conducted a total of 40 replications – 10 for each climatic context (slow and fast) and plasticity cost (without and with). Prior to running the final simulations, we conducted investigative simulations to test assumptions and verify rate stabilization. At the end of each simulation, species, individuals, phylogeny, and other variables were saved based on events that occurred in the world. We also saved the calculated rates, including (1) trait evolution, which corresponds to the mean difference in internal temperature between the time t and t-1, divided by the trait value in t-1; (2) speciation rate, which corresponds to the number of speciation events in the time t, divided by the number of species in the time t-1; (3) extinction rate, which corresponds to the number of extinction events in time t, divided by the number of species in time t-1; and (4) diversification rate, which corresponds to the difference between the speciation and extinction rates.

### ANALYSIS

We used an Analysis of Variance (ANOVA) in R to verify the effect of plasticity on trait evolution, speciation, extinction, and diversification rates under different climatic contexts and different plasticity costs. To identify which contexts differ from one another, we conducted a Tukey’s test. Additionally, we checked for differences in abundance and occupancy among plasticity levels across climatic contexts and different plasticity costs.

## Results

We found the same pattern of results for plasticity in climatic contexts and plasticity costs, and all results have a p-value < 2x10^-16^, and F (7, 72) > 58. An increase in plasticity leads to a decrease in extinction and trait evolution rates. Systems with species exhibiting plasticity also experience higher speciation rates. Additionally, as plasticity increases in systems, diversification rates also increase. In all plots, higher levels of plasticity have a smaller standard deviation, indicating that the results consistently converge to a single outcome (Figure 1). Furthermore, our analysis using Tukey’s test revealed significant differences in almost all pairwise comparisons among the factor levels (found in Tables 1, 2, 3, and 4 in the Supplementary Material).

**Figure 1:**
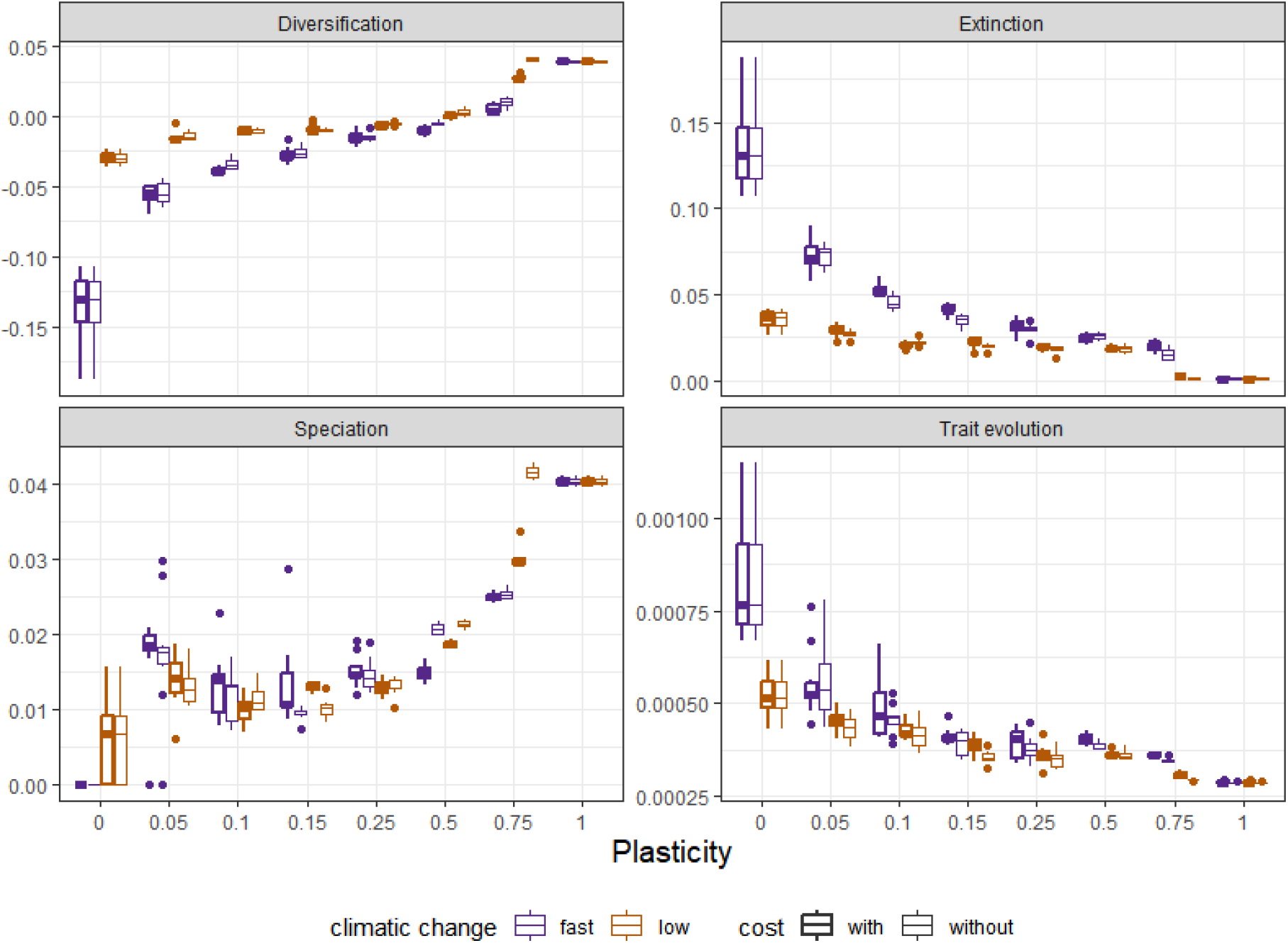
Effect of plasticity on adaptive evolution in the contexts of fast and slow climatic change, and in the presence or absence of plasticity costs. (1) effect of the plasticity on diversification rate; (2) effect of the plasticity on extinction rate; (3) effect of the plasticity on speciation rate; and (4) effect of the plasticity on trait evolution rate. The X scale was designed to cover plasticity differences from 0 to 1 with a progression of 0.25. Additional factor levels between 0 and 0.25 were included to examine patterns in the abrupt change between these factor levels.

Systems with plasticity equal to or less than 0.5 experienced the extinction of all species after the initial organization of the world, with a minimum threshold of 60 timesteps. The results presented in the graphs display outcomes before the mass extinction, except for cases with plasticity values of 0.75 and 1, which reached 50,000 species – high plasticity postpones mass extinctions events. In general, we observed higher rates of extinction and trait evolution in the context of rapid climatic changes. Speciation and diversification showed similar results in both climatic contexts, but the threshold for achieving positive diversification was higher in the case of rapid climatic changes. The plasticity cost context yielded similar results in both climatic change scenarios (p-value > 0.49 and F (1, 318) < 0.47), but in the slow climatic change context with plasticity cost, maximum plasticity was required to achieve higher speciation and diversification rates. Additionally, we observed that as plasticity increases, both occupancy and abundance also increase (Figure 2). Abundance and occupancy are almost always higher in simulations without plasticity costs compared to systems with plasticity costs (Figure 2).

**Figure 2:**
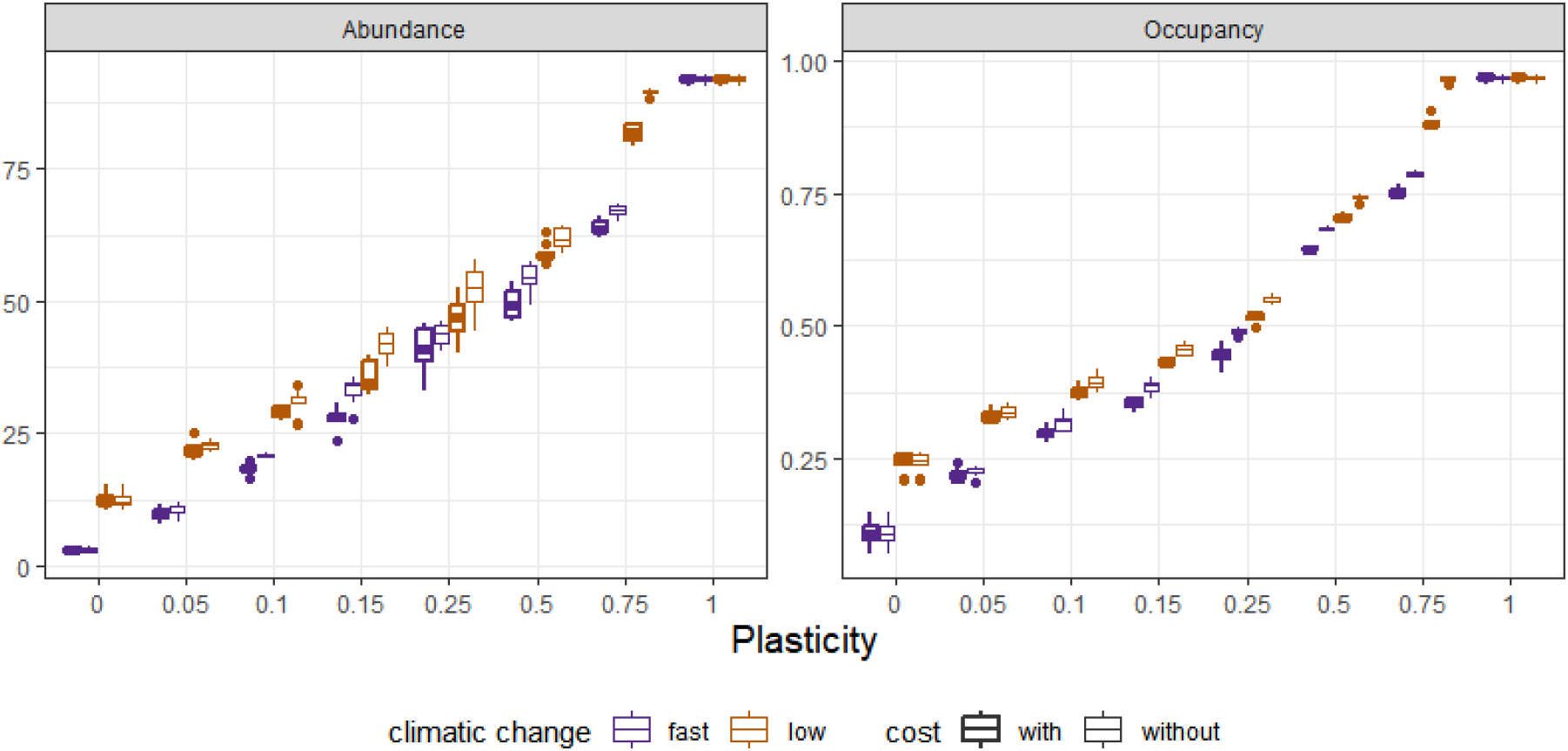
Effect of plasticity on abundance and occupancy in context of fast and slow climatic change and presence and absence of the plasticity cost – mechanisms. The X scale was designed to cover plasticity differences from 0 to 1 with a progression of 0.25. Additional factor levels between 0 and 0.25 were included to examine patterns in the abrupt change between these factor levels.

## Discussion

We found that as plasticity increases, there is a decrease in extinction and trait evolution rates in both climatic contexts. Systems with individuals having plasticity have higher speciation rates. Additionally, as plasticity increases, diversification rates also increase. The observed increase in speciation and diversification rates in systems with plasticity supports the hypothesis that plasticity facilitates the adaptive evolution of species in response to environmental changes (Ghalambor et al., 2015; Wong and Candolin, 2015; Fox et al., 2019; Pfennig, 2021).

The support for both facilitation hypotheses (“Plasticity-First” and “Buying Time” hypotheses) can be explained by several factors. Low plasticity, in comparison with high plasticity, would lead to the survival of only a few individuals, low world occupancy, and significant distances between individuals (decreasing genetic exchanges). In this scenario, a small population can establish itself as minimally viable, accumulate genetic differences rapidly, and undergo speciation (in comparison to systems without plasticity) (Baldwin, 1896). Conversely, increases in plasticity lead to larger population sizes, higher spatialized world occupancy, increased genetic exchanges, reduced free spaces, and formation of groups of subpopulations with accumulated differences, thus favoring speciation. These findings emphasize the complex and dynamic relationship between plasticity and adaptive evolution. Additionally, they indicate relevant factors and mechanisms, such as the nature of environmental changes, isolation, population size, occupancy or niche, capacity to form groups, plasticity extension, and dispersal capacity.

The finding that increased plasticity is associated with decreased extinction rates suggests that plasticity can act as a buffer against environmental fluctuations, allowing species to adapt to changing conditions and avoid extinction through evolutionary rescue (recovery and persistence of a population through natural selection) (Bell, 2017). This is consistent with previous studies showing that plasticity can increase the fitness of individuals and enhance their ability to cope with environmental stress (Scheiner et al., 2020; Barbosa-Júnior et al., 2022) and facilitate adaptation (Scheiner et al., 2020). In our case, we support the results regarding extinction at an macroevolutionary scale. Although low plasticity can lead to a minimum viable population and thus decrease extinction, it may sometimes also be associated with a specific habitat or ecological niche (Futuyama and Moreno, 1988). This can make species more vulnerable to extinction if their habitat or niche disappears; specialist species, in particular, can be more prone to extinction (Raia et al., 2016). Our results show that in both moderate and extreme environmental changes, species with low plasticity did not survive and became extinct. Only species with high plasticity were viable in such situations, and the extinction was significantly higher and faster in cases of extreme changes.

The relevance and current interest in plasticity cost have grown due to its potential influence on adaptive evolution (Snell-Rood and Ehlman, 2021). Plasticity is widespread in nature as an adaptive trait (Pfenning, 2021), but the associated costs could potentially limit the level of adaptation (DeWitt et al., 1998). In our simulations, in both climatic change scenarios, the impact of plasticity cost on the outcomes was similar. The absence of a significant difference in plasticity cost arises from the necessity to use a sufficiently low plasticity cost to run simulations without experiencing mass extinction during the initial phase, allowing us to obtain comparable results. In another hypothetical scenario, the slight differences observed across climatic contexts, in terms of speciation, diversification, abundance and occupancy, would likely be more pronounced. One possibility is that costlier plasticity would require higher levels of plasticity to achieve positive diversification and maintain viable populations. In such cases, plasticity cost can increase extinction rates and decrease diversification. Our observed differences and the rapid convergence to mass extinction with minimal plasticity costs provide further support for this notion.

We found that as plasticity increases, systems exhibit decreased trait evolution rates. This can be explained by the observation that systems with high plasticity do not necessarily need to evolve traits since traits can adjust in the direction of the optimal trait. In contrast, systems with little or no plasticity experience major changes in traits primarily through random mutations, resulting in more significant trait evolution among generations. Our work did not test completely the ‘Plasticity-First’ hypothesis because we would need to enhance the complexity of our model to include genetic assimilation. But through a simpler model, we were able to discover relevant mechanisms and effectively control the parameter space, as suggested by Pfenning et al. (2010), which facilitated establishing causality. We recommend conducting additional empirical studies to validate the effects of the mentioned mechanisms, such as population size, group formation, gene exchange, and plasticity extension.

## Conclusion

Our models indicate that higher plasticity may allow organisms to better cope with environmental fluctuations and avoid extinction. In contrast, low plasticity may promote rapid speciation compared to systems without plasticity, but it may also increase the risk of extinction, especially in extreme climatic contexts. Additionally, higher plasticity increases speciation and diversification rates in both climatic contexts, supporting the facilitation hypothesis. Our study provides valuable insights into understanding the role of plasticity in adaptive evolution, emphasizing the delicate balance in facilitating evolutionary processes. Factors such as the nature of environmental changes, isolation, population size, niche occupancy, group formation capacity, plasticity extension, and dispersal capacity all contribute to the complex dynamics between plasticity and adaptive evolution. These results highlight the importance of considering plasticity in studies of evolutionary dynamics and offer a new understanding of the effect of plasticity on adaptive evolution at macroevolutionary scales, particularly in the context of human-induced rapid environmental changes (Wong and Candolin, 2015; Bro-Jorgensen et al., 2019). By doing so, we aim to provide greater clarity about the mechanisms and influence of plasticity on adaptive evolution, helping to bridge the gap in the literature, the theory of evolution, and the new extended synthesis (Laland, 2015; Scheiner and Mindell, 2020).

## Acknowledgements

We are grateful to Fundação de Amparo à Pesquisa do Estado da Bahia (FAPESB) for awarding the PhD scholarship (BOL0207/2020) to ECBJ. We also thank Dr. Henrique Batalha Filho for his valuable comments and suggestions on the work.

