## Supplementary Material for "The effects of phenotypic plasticity on diversification rates and adaptive evolution in simulated environments with different climatic and cost contexts"

### SUPLEMENTARY MATERIAL

#### Detailed methods:

We set a spatial cost of 0.001 for land and 0.002 for water (the default), and each cell had eight neighbors. The temperature range was from 0 to 1. In the slow climatic context, temperature values took 1000 timesteps to transition from one extreme temperature (colder) to the other extreme (warmer) – from 0 to 1 in increments of 0.001 for each time step. On the other hand, in fast climate changes, the temperature changed twice as fast as in the slow context, taking half the time to transition from one extreme value to another, with changes of 0.002 in each time step. We also introduced spatial temperature variations at all timesteps, which were based on latitudinal and longitudinal variations observed in the real world, with warmer temperatures in the center and the coldest temperatures farther away (there was a little longitudinal variation). Although our simulation used a temperature scale of 0 to 1, we patterned it after real-world values by converting Celsius degrees to a 0-1 scale.

This was accomplished based on modelling temperature patterns: 0.01-1 (100 amplitude values), with the center value being 0.5. Real-world temperature follows a pattern of 0-30; 60 amplitude values, with the center value being 30. In a range of 100 amplitude values, 0.6 corresponds to 1 value within the 0-30 real temperature range. Our map covers 0 latitude to 40 latitude. In the temperature world map, this corresponds to 30 to 10 degrees Celsius (a 20-degree difference – 66%). In modelling temperature, 0.5 * 0.6 (66%) = ~0.35 both below and above, so the modelling temperature has a maximum of 0.85 and a minimum of 0.15. We have 800 latitude cells, with cell 400 representing 0 latitude. Therefore: 0 Latitude cell = 10 Celsius degrees = 0.15 modelling temperature; 400 Latitude cell = 30 Celsius degrees = 0.5 modelling temperature; and 800 Latitude cell = 10 Celsius degrees = 0.85 modelling temperature (more details are provided in the Supplementary Material). To calculate the progression from the 400th latitude cell to the extreme values, we divide the amplitude value of 0.35 by 399 cell numbers, resulting in 0.00087. We then increment/decrement the central cell by 0.00087 modelling temperature values from the 400th latitude cell to the 0th and 800th cells, reaching the extreme values of 0.15 and 0.85, respectively.

**Table 1: Tukey's test results correspond to simulations that verify the effect of phenotypic plasticity on adaptive rates in the context of slow climatic changes and absence of the plasticity cost**

| | | | 95% Confidence Interval | | |
|---|---|---|---|---|---|
| **Variables** | Factor levels | Mean difference | Lower bound | Upper bound | Significance |
| **Trait evolution** | 0.05-0 | -0.00009 | -0.0001307 | -0.0000498 | 0.00000 |
| | 0.1-0 | -0.0001059 | -0.0001464 | -0.0000655 | 0.00000 |
| | 0.15-0 | -0.0001677 | -0.0002081 | -0.0001272 | 0.00000 |
| | 0.25-0 | -0.0001740 | -0.0002145 | -0.0001335 | 0.00000 |
| | 0.5-0 | -0.0001626 | -0.0002030 | -0.0001221 | 0.00000 |
| | 0.75-0 | -0.0002295 | -0.0002699 | -0.0001890 | 0.00000 |
| | 1-0 | -0.0002379 | -0.0002784 | -0.0001975 | 0.00000 |
| | 0.1-0.05 | -0.0000157 | -0.0000562 | 0.0000248 | 0.92623 |
| | 0.15-0.05 | -0.0000774 | -0.0001178 | -0.0000369 | 0.00000 |
| | 0.25-0.05 | -0.0000837 | -0.0001242 | -0.0000433 | 0.00000 |
| | 0.5-0.05 | -0.0000723 | -0.0001127 | -0.0000318 | 0.00001 |
| | 0.75-0.05 | -0.0001392 | -0.0001797 | -0.0000987 | 0.00000 |
| | 1-0.05 | -0.0001476 | -0.0001881 | -0.0001072 | 0.00000 |
| | 0.15-0.1 | -0.0000617 | -0.0001022 | -0.0000212 | 0.00025 |
| | 0.25-0.1 | -0.0000680 | -0.0001085 | -0.0000276 | 0.00004 |
| | 0.5-0.1 | -0.0000566 | -0.0000970 | -0.0000161 | 0.00104 |
| | 0.75-0.1 | -0.0001235 | -0.0001639 | -0.0000830 | 0.00000 |
| | 1-0.1 | -0.0001319 | -0.0001724 | -0.0000915 | 0.00000 |
| | 0.25-0.15 | -0.0000063 | -0.0000468 | 0.0000341 | 0.99968 |
| | 0.5-0.15 | 0.0000051 | -0.0000354 | 0.0000456 | 0.99992 |
| | 0.75-0.15 | -0.0000618 | -0.0001023 | -0.0000213 | 0.00024 |
| | 1-0.15 | -0.0000703 | -0.0001107 | -0.0000298 | 0.00002 |
| | 0.5-0.25 | 0.0000114 | -0.0000290 | 0.0000519 | 0.98673 |
| | 0.75-0.25 | -0.0000555 | -0.0000959 | -0.0000150 | 0.00140 |
| | 1-0.25 | -0.0000639 | -0.0001044 | -0.0000235 | 0.00013 |
| | 0.75-0.5 | -0.0000669 | -0.0001074 | -0.0000265 | 0.00005 |
| | 1-0.5 | -0.0000754 | -0.0001158 | -0.0000349 | 0.00000 |
| | 1-0.75 | -0.0000084 | -0.0000489 | 0.0000320 | 0.99795 |

| | | | | | |
|---|---|---|---|---|---|
| **Speciation** | 0.05-0 | 0.00732 | 0.00406 | 0.01057 | 0.00000 |
| | 0.1-0 | 0.00591 | 0.00266 | 0.00917 | 0.00001 |
| | 0.15-0 | 0.00454 | 0.00129 | 0.00780 | 0.00108 |
| | 0.25-0 | 0.00760 | 0.00434 | 0.01085 | 0.00000 |
| | 0.5-0 | 0.01566 | 0.01241 | 0.01892 | 0.00000 |
| | 0.75-0 | 0.03602 | 0.03277 | 0.03928 | 0.00000 |
| | 1-0 | 0.03475 | 0.03150 | 0.03801 | 0.00000 |
| | 0.1-0.05 | -0.00140 | -0.00466 | 0.00185 | 0.87855 |
| | 0.15-0.05 | -0.00277 | -0.00603 | 0.00048 | 0.15258 |
| | 0.25-0.05 | 0.00028 | -0.00297 | 0.00354 | 0.99999 |
| | 0.5-0.05 | 0.00835 | 0.00509 | 0.01160 | 0.00000 |
| | 0.75-0.05 | 0.02871 | 0.02545 | 0.03196 | 0.00000 |
| | 1-0.05 | 0.02744 | 0.02418 | 0.03069 | 0.00000 |
| | 0.15-0.1 | -0.00137 | -0.00463 | 0.00188 | 0.89038 |
| | 0.25-0.1 | 0.00168 | -0.00157 | 0.00493 | 0.74002 |
| | 0.5-0.1 | 0.00974 | 0.00649 | 0.01300 | 0.00000 |
| | 0.75-0.1 | 0.03010 | 0.02685 | 0.03336 | 0.00000 |
| | 1-0.1 | 0.02884 | 0.02558 | 0.03209 | 0.00000 |
| | 0.25-0.15 | 0.00305 | -0.00020 | 0.00631 | 0.08164 |
| | 0.5-0.15 | 0.01112 | 0.00786 | 0.01438 | 0.00000 |
| | 0.75-0.15 | 0.03148 | 0.02822 | 0.03473 | 0.00000 |
| | 1-0.15 | 0.03021 | 0.02695 | 0.03346 | 0.00000 |
| | 0.5-0.25 | 0.00807 | 0.00481 | 0.01132 | 0.00000 |
| | 0.75-0.25 | 0.02842 | 0.02517 | 0.03168 | 0.00000 |
| | 1-0.25 | 0.02715 | 0.02390 | 0.03041 | 0.00000 |
| | 0.75-0.5 | 0.02036 | 0.01710 | 0.02361 | 0.00000 |
| | 1-0.5 | 0.01908 | 0.01583 | 0.02234 | 0.00000 |
| | 1-0.75 | -0.00127 | -0.00453 | 0.00199 | 0.92394 |

| | | | | | |
|---|---|---|---|---|---|
| **Extinction** | 0.05-0 | -0.0085 | -0.0117 | -0.0052 | 0.0000 |
| | 0.1-0 | -0.0131 | -0.0164 | -0.0098 | 0.0000 |
| | 0.15-0 | -0.0155 | -0.0187 | -0.0122 | 0.0000 |
| | 0.25-0 | -0.0170 | -0.0203 | -0.0137 | 0.0000 |
| | 0.5-0 | -0.0170 | -0.0203 | -0.0137 | 0.0000 |
| | 0.75-0 | -0.0344 | -0.0376 | -0.0311 | 0.0000 |
| | 1-0 | -0.0344 | -0.0377 | -0.0311 | 0.0000 |
| | 0.1-0.05 | -0.0046 | -0.0079 | -0.0014 | 0.0009 |
| | 0.15-0.05 | -0.0070 | -0.0103 | -0.0037 | 0.0000 |
| | 0.25-0.05 | -0.0086 | -0.0118 | -0.0053 | 0.0000 |
| | 0.5-0.05 | -0.0085 | -0.0118 | -0.0053 | 0.0000 |
| | 0.75-0.05 | -0.0259 | -0.0292 | -0.0226 | 0.0000 |
| | 1-0.05 | -0.0259 | -0.0292 | -0.0226 | 0.0000 |
| | 0.15-0.1 | -0.0024 | -0.0057 | 0.0009 | 0.3319 |
| | 0.25-0.1 | -0.0039 | -0.0072 | -0.0006 | 0.0087 |
| | 0.5-0.1 | -0.0039 | -0.0072 | -0.0006 | 0.0089 |
| | 0.75-0.1 | -0.0213 | -0.0245 | -0.0180 | 0.0000 |
| | 1-0.1 | -0.0213 | -0.0246 | -0.0180 | 0.0000 |
| | 0.25-0.15 | -0.0015 | -0.0048 | 0.0017 | 0.8200 |
| | 0.5-0.15 | -0.0015 | -0.0048 | 0.0017 | 0.8238 |
| | 0.75-0.15 | -0.0189 | -0.0222 | -0.0156 | 0.0000 |
| | 1-0.15 | -0.0189 | -0.0222 | -0.0156 | 0.0000 |
| | 0.5-0.25 | 0.0000 | -0.0033 | 0.0033 | 1.0000 |
| | 0.75-0.25 | -0.0173 | -0.0206 | -0.0141 | 0.0000 |
| | 1-0.25 | -0.0174 | -0.0206 | -0.0141 | 0.0000 |
| | 0.75-0.5 | -0.0174 | -0.0206 | -0.0141 | 0.0000 |
| | 1-0.5 | -0.0174 | -0.0206 | -0.0141 | 0.0000 |
| | 1-0.75 | 0.0000 | -0.0033 | 0.0033 | 1.0000 |

| | | | | | |
|---|---|---|---|---|---|
| **Diversification** | 0.05-0 | 0.01577 | 0.01278 | 0.01876 | 0.00000 |
| | 0.1-0 | 0.01901 | 0.01602 | 0.02199 | 0.00000 |
| | 0.15-0 | 0.02001 | 0.01702 | 0.02299 | 0.00000 |
| | 0.25-0 | 0.02461 | 0.02162 | 0.02759 | 0.00000 |
| | 0.5-0 | 0.03267 | 0.02968 | 0.03566 | 0.00000 |
| | 0.75-0 | 0.07038 | 0.06739 | 0.07337 | 0.00000 |
| | 1-0 | 0.06912 | 0.06613 | 0.07211 | 0.00000 |
| | 0.1-0.05 | 0.00324 | 0.00025 | 0.00623 | 0.02468 |
| | 0.15-0.05 | 0.00424 | 0.00125 | 0.00723 | 0.00085 |
| | 0.25-0.05 | 0.00883 | 0.00585 | 0.01183 | 0.00000 |
| | 0.5-0.05 | 0.01689 | 0.01391 | 0.01989 | 0.00000 |
| | 0.75-0.05 | 0.05461 | 0.05162 | 0.05760 | 0.00000 |
| | 1-0.05 | 0.05335 | 0.05036 | 0.05634 | 0.00000 |
| | 0.15-0.1 | 0.00100 | -0.00199 | 0.00399 | 0.96546 |
| | 0.25-0.1 | 0.00560 | 0.00261 | 0.00859 | 0.00000 |
| | 0.5-0.1 | 0.01366 | 0.01067 | 0.01665 | 0.00000 |
| | 0.75-0.1 | 0.05138 | 0.04839 | 0.05436 | 0.00000 |
| | 1-0.1 | 0.05011 | 0.04712 | 0.05310 | 0.00000 |
| | 0.25-0.15 | 0.00460 | 0.00161 | 0.00759 | 0.00021 |
| | 0.5-0.15 | 0.01266 | 0.00967 | 0.01564 | 0.00000 |
| | 0.75-0.15 | 0.05037 | 0.04738 | 0.05336 | 0.00000 |
| | 1-0.15 | 0.04911 | 0.04612 | 0.05210 | 0.00000 |
| | 0.5-0.25 | 0.00805 | 0.00507 | 0.01104 | 0.00000 |
| | 0.75-0.25 | 0.04577 | 0.04278 | 0.04876 | 0.00000 |
| | 1-0.25 | 0.04451 | 0.04152 | 0.04750 | 0.00000 |
| | 0.75-0.5 | 0.03771 | 0.03473 | 0.04070 | 0.00000 |
| | 1-0.5 | 0.03645 | 0.03346 | 0.03944 | 0.00000 |
| | 1-0.75 | -0.00126 | -0.00425 | 0.00172 | 0.88929 |

**Table 2: Tukey's test results correspond to simulations that verify the effect of phenotypic plasticity on adaptive rates in the context of slow climatic changes and presence of the plasticity cost**

| | | | 95% Confidence Interval | | |
|---|---|---|---|---|---|
| **Variables** | Factor levels | Mean difference | Lower bound | Upper bound | Significance |

| | | | | | |
|---|---|---|---|---|---|
| **Trait evolution** | 0.05-0 | -0.00007 | -0.00011 | -0.00003 | 0.00000 |
| | 0.1-0 | -0.00010 | -0.00014 | -0.00006 | 0.00000 |
| | 0.15-0 | -0.00013 | -0.00017 | -0.00010 | 0.00000 |
| | 0.25-0 | -0.00017 | -0.00020 | -0.00013 | 0.00000 |
| | 0.5-0 | -0.00016 | -0.00020 | -0.00012 | 0.00000 |
| | 0.75-0 | -0.00022 | -0.00026 | -0.00018 | 0.00000 |
| | 1-0 | -0.00024 | -0.00027 | -0.00020 | 0.00000 |
| | 0.1-0.05 | -0.00003 | -0.00006 | 0.00001 | 0.44062 |
| | 0.15-0.05 | -0.00006 | -0.00010 | -0.00002 | 0.00008 |
| | 0.25-0.05 | -0.00009 | -0.00013 | -0.00006 | 0.00000 |
| | 0.5-0.05 | -0.00009 | -0.00013 | -0.00005 | 0.00000 |
| | 0.75-0.05 | -0.00014 | -0.00018 | -0.00011 | 0.00000 |
| | 1-0.05 | -0.00016 | -0.00020 | -0.00013 | 0.00000 |
| | 0.15-0.1 | -0.00004 | -0.00007 | 0.00000 | 0.07089 |
| | 0.25-0.1 | -0.00007 | -0.00011 | -0.00003 | 0.00001 |
| | 0.5-0.1 | -0.00006 | -0.00010 | -0.00002 | 0.00007 |
| | 0.75-0.1 | -0.00012 | -0.00016 | -0.00008 | 0.00000 |
| | 1-0.1 | -0.00014 | -0.00018 | -0.00010 | 0.00000 |
| | 0.25-0.15 | -0.00003 | -0.00007 | 0.00001 | 0.17609 |
| | 0.5-0.15 | -0.00003 | -0.00006 | 0.00001 | 0.41353 |
| | 0.75-0.15 | -0.00008 | -0.00012 | -0.00004 | 0.00000 |
| | 1-0.15 | -0.00010 | -0.00014 | -0.00006 | 0.00000 |
| | 0.5-0.25 | 0.00001 | -0.00003 | 0.00004 | 0.99974 |
| | 0.75-0.25 | -0.00005 | -0.00009 | -0.00001 | 0.00185 |
| | 1-0.25 | -0.00007 | -0.00011 | -0.00003 | 0.00000 |
| | 0.75-0.5 | -0.00006 | -0.00010 | -0.00002 | 0.00035 |
| | 1-0.5 | -0.00008 | -0.00011 | -0.00004 | 0.00000 |
| | 1-0.75 | -0.00002 | -0.00006 | 0.00002 | 0.76428 |

| | | | | | |
|---|---|---|---|---|---|
| **Speciation** | 0.05-0 | 0.00834 | 0.00486 | 0.01183 | 0.00000 |
| | 0.1-0 | 0.00420 | 0.00072 | 0.00768 | 0.00768 |
| | 0.15-0 | 0.00745 | 0.00397 | 0.01093 | 0.00000 |
| | 0.25-0 | 0.00727 | 0.00378 | 0.01075 | 0.00000 |
| | 0.5-0 | 0.01306 | 0.00957 | 0.01654 | 0.00000 |
| | 0.75-0 | 0.02443 | 0.02095 | 0.02792 | 0.00000 |
| | 1-0 | 0.03475 | 0.03127 | 0.03823 | 0.00000 |
| | 0.1-0.05 | -0.00414 | -0.00762 | -0.00066 | 0.00902 |
| | 0.15-0.05 | -0.00089 | -0.00437 | 0.00258 | 0.99248 |
| | 0.25-0.05 | -0.00108 | -0.00456 | 0.00240 | 0.97776 |
| | 0.5-0.05 | 0.00471 | 0.00123 | 0.00819 | 0.00169 |
| | 0.75-0.05 | 0.01609 | 0.01261 | 0.01957 | 0.00000 |
| | 1-0.05 | 0.02640 | 0.02292 | 0.02988 | 0.00000 |
| | 0.15-0.1 | 0.00324 | -0.00023 | 0.00673 | 0.08498 |
| | 0.25-0.1 | 0.00307 | -0.00041 | 0.00654 | 0.12489 |
| | 0.5-0.1 | 0.00885 | 0.00537 | 0.01233 | 0.00000 |
| | 0.75-0.1 | 0.02023 | 0.01675 | 0.02372 | 0.00000 |
| | 1-0.1 | 0.03055 | 0.02707 | 0.03403 | 0.00000 |
| | 0.25-0.15 | -0.00018 | -0.00366 | 0.00329 | 1.00000 |
| | 0.5-0.15 | 0.00560 | 0.00212 | 0.00908 | 0.00009 |
| | 0.75-0.15 | 0.01698 | 0.01350 | 0.02047 | 0.00000 |
| | 1-0.15 | 0.02730 | 0.02382 | 0.03078 | 0.00000 |
| | 0.5-0.25 | 0.00578 | 0.00230 | 0.00927 | 0.00005 |
| | 0.75-0.25 | 0.01717 | 0.01369 | 0.02065 | 0.00000 |
| | 1-0.25 | 0.02748 | 0.02400 | 0.03096 | 0.00000 |
| | 0.75-0.5 | 0.01138 | 0.00790 | 0.01486 | 0.00000 |
| | 1-0.5 | 0.02170 | 0.01821 | 0.02518 | 0.00000 |
| | 1-0.75 | 0.01031 | 0.00683 | 0.01379 | 0.00000 |

| | | | | | |
|---|---|---|---|---|---|
| **Extinction** | 0.05-0 | -0.00630 | -0.00998 | -0.00261 | 0.00003 |
| | 0.1-0 | -0.01487 | -0.01855 | -0.01118 | 0.00000 |
| | 0.15-0 | -0.01363 | -0.01732 | -0.00994 | 0.00000 |
| | 0.25-0 | -0.01609 | -0.01978 | -0.01240 | 0.00000 |
| | 0.5-0 | -0.01690 | -0.02059 | -0.01322 | 0.00000 |
| | 0.75-0 | -0.03290 | -0.03658 | -0.02921 | 0.00000 |
| | 1-0 | -0.03437 | -0.03806 | -0.03068 | 0.00000 |
| | 0.1-0.05 | -0.00857 | -0.01226 | -0.00488 | 0.00000 |
| | 0.15-0.05 | -0.00733 | -0.01102 | -0.00365 | 0.00000 |
| | 0.25-0.05 | -0.00980 | -0.01348 | -0.00611 | 0.00000 |
| | 0.5-0.05 | -0.01061 | -0.01429 | -0.00692 | 0.00000 |
| | 0.75-0.05 | -0.02660 | -0.03029 | -0.02291 | 0.00000 |
| | 1-0.05 | -0.02807 | -0.03176 | -0.02439 | 0.00000 |
| | 0.15-0.1 | 0.00124 | -0.00245 | 0.00492 | 0.96525 |
| | 0.25-0.1 | -0.00123 | -0.00491 | 0.00246 | 0.96701 |
| | 0.5-0.1 | -0.00204 | -0.00572 | 0.00165 | 0.67211 |
| | 0.75-0.1 | -0.01803 | -0.02172 | -0.01434 | 0.00000 |
| | 1-0.1 | -0.01950 | -0.02319 | -0.01581 | 0.00000 |
| | 0.25-0.15 | -0.00246 | -0.00615 | 0.00123 | 0.43470 |
| | 0.5-0.15 | -0.00327 | -0.00696 | 0.00041 | 0.11890 |
| | 0.75-0.15 | -0.01927 | -0.02295 | -0.01558 | 0.00000 |
| | 1-0.15 | -0.02074 | -0.02443 | -0.01705 | 0.00000 |
| | 0.5-0.25 | -0.00081 | -0.00450 | 0.00288 | 0.99713 |
| | 0.75-0.25 | -0.01680 | -0.02049 | -0.01312 | 0.00000 |
| | 1-0.25 | -0.01828 | -0.02196 | -0.01459 | 0.00000 |
| | 0.75-0.5 | -0.01599 | -0.01968 | -0.01231 | 0.00000 |
| | 1-0.5 | -0.01747 | -0.02115 | -0.01378 | 0.00000 |
| | 1-0.75 | -0.00147 | -0.00516 | 0.00221 | 0.91473 |

| | | | | | |
|---|---|---|---|---|---|
| **Diversification** | 0.05-0 | 0.01464 | 0.01104 | 0.01824 | 0.00000 |
| | 0.1-0 | 0.01907 | 0.01547 | 0.02266 | 0.00000 |
| | 0.15-0 | 0.02108 | 0.01748 | 0.02468 | 0.00000 |
| | 0.25-0 | 0.02336 | 0.01976 | 0.02695 | 0.00000 |
| | 0.5-0 | 0.02996 | 0.02636 | 0.03355 | 0.00000 |
| | 0.75-0 | 0.05734 | 0.05374 | 0.06093 | 0.00000 |
| | 1-0 | 0.06912 | 0.06552 | 0.07271 | 0.00000 |
| | 0.1-0.05 | 0.00443 | 0.00083 | 0.00802 | 0.00603 |
| | 0.15-0.05 | 0.00644 | 0.00284 | 0.01004 | 0.00001 |
| | 0.25-0.05 | 0.00872 | 0.00512 | 0.01231 | 0.00000 |
| | 0.5-0.05 | 0.01532 | 0.01172 | 0.01891 | 0.00000 |
| | 0.75-0.05 | 0.04270 | 0.03910 | 0.04629 | 0.00000 |
| | 1-0.05 | 0.05448 | 0.05088 | 0.05807 | 0.00000 |
| | 0.15-0.1 | 0.00201 | -0.00158 | 0.00561 | 0.65736 |
| | 0.25-0.1 | 0.00429 | 0.00069 | 0.00789 | 0.00876 |
| | 0.5-0.1 | 0.01089 | 0.00729 | 0.01449 | 0.00000 |
| | 0.75-0.1 | 0.03827 | 0.03467 | 0.04186 | 0.00000 |
| | 1-0.1 | 0.05005 | 0.04646 | 0.05365 | 0.00000 |
| | 0.25-0.15 | 0.00228 | -0.00132 | 0.00588 | 0.50336 |
| | 0.5-0.15 | 0.00888 | 0.00528 | 0.01248 | 0.00000 |
| | 0.75-0.15 | 0.03626 | 0.03266 | 0.03985 | 0.00000 |
| | 1-0.15 | 0.04804 | 0.04445 | 0.05164 | 0.00000 |
| | 0.5-0.25 | 0.00660 | 0.00300 | 0.01020 | 0.00001 |
| | 0.75-0.25 | 0.03398 | 0.03038 | 0.03757 | 0.00000 |
| | 1-0.25 | 0.04576 | 0.04217 | 0.04936 | 0.00000 |
| | 0.75-0.5 | 0.02738 | 0.02378 | 0.03097 | 0.00000 |
| | 1-0.5 | 0.03916 | 0.03557 | 0.04276 | 0.00000 |
| | 1-0.75 | 0.01179 | 0.00819 | 0.01538 | 0.00000 |

**Table 3: Tukey's test results correspond to simulations that verify the effect of phenotypic plasticity on adaptive rates in the context of fast climatic changes and absence of the plasticity cost**

| | | | 95% Confidence Interval | | |
|---|---|---|---|---|---|
| **Variables** | Factor levels | Mean difference | Lower bound | Upper bound | Significance |

| | | | | | |
|---|---|---|---|---|---|
| **Trait evolution** | 0.05-0 | -0.00027 | -0.00037 | -0.00017 | 0.00000 |
| | 0.1-0 | -0.00037 | -0.00047 | -0.00027 | 0.00000 |
| | 0.15-0 | -0.00043 | -0.00053 | -0.00034 | 0.00000 |
| | 0.25-0 | -0.00045 | -0.00055 | -0.00035 | 0.00000 |
| | 0.5-0 | -0.00044 | -0.00054 | -0.00034 | 0.00000 |
| | 0.75-0 | -0.00048 | -0.00058 | -0.00038 | 0.00000 |
| | 1-0 | -0.00054 | -0.00064 | -0.00044 | 0.00000 |
| | 0.1-0.05 | -0.00010 | -0.00020 | -0.00000 | 0.04262 |
| | 0.15-0.05 | -0.00017 | -0.00026 | -0.00007 | 0.00004 |
| | 0.25-0.05 | -0.00018 | -0.00028 | -0.00008 | 0.00001 |
| | 0.5-0.05 | -0.00017 | -0.00027 | -0.00007 | 0.00002 |
| | 0.75-0.05 | -0.00021 | -0.00031 | -0.00011 | 0.00000 |
| | 1-0.05 | -0.00027 | -0.00037 | -0.00017 | 0.00000 |
| | 0.15-0.1 | -0.00007 | -0.00016 | 0.00003 | 0.44498 |
| | 0.25-0.1 | -0.00008 | -0.00018 | 0.00002 | 0.18043 |
| | 0.5-0.1 | -0.00007 | -0.00017 | 0.00003 | 0.33754 |
| | 0.75-0.1 | -0.00011 | -0.00021 | -0.00001 | 0.01558 |
| | 1-0.1 | -0.00017 | -0.00027 | -0.00007 | 0.00002 |
| | 0.25-0.15 | -0.00002 | -0.00011 | 0.00008 | 0.99957 |
| | 0.5-0.15 | -0.00001 | -0.00010 | 0.00009 | 0.99999 |
| | 0.75-0.15 | -0.00005 | -0.00014 | 0.00005 | 0.82167 |
| | 1-0.15 | -0.00011 | -0.00021 | -0.00001 | 0.02292 |
| | 0.5-0.25 | 0.00001 | -0.00009 | 0.00011 | 0.99997 |
| | 0.75-0.25 | -0.00003 | -0.00013 | 0.00007 | 0.97901 |
| | 1-0.25 | -0.00009 | -0.00019 | 0.00001 | 0.08927 |
| | 0.75-0.5 | -0.00004 | -0.00014 | 0.00006 | 0.89976 |
| | 1-0.5 | -0.00010 | -0.00020 | -0.00000 | 0.03778 |
| | 1-0.75 | -0.00006 | -0.00016 | 0.00004 | 0.53112 |

| | | | | | |
|---|---|---|---|---|---|
| **Speciation** | 0.05-0 | 0.01738 | 0.01286 | 0.02190 | 0.00000 |
| | 0.1-0 | 0.01167 | 0.00715 | 0.01619 | 0.00000 |
| | 0.15-0 | 0.00940 | 0.00488 | 0.01392 | 0.00000 |
| | 0.25-0 | 0.01454 | 0.01002 | 0.01906 | 0.00000 |
| | 0.5-0 | 0.02072 | 0.01620 | 0.02525 | 0.00000 |
| | 0.75-0 | 0.02535 | 0.02083 | 0.02988 | 0.00000 |
| | 1-0 | 0.04041 | 0.03589 | 0.04494 | 0.00000 |
| | 0.1-0.05 | -0.00571 | -0.01023 | -0.00118 | 0.00442 |
| | 0.15-0.05 | -0.00798 | -0.01250 | -0.00346 | 0.00001 |
| | 0.25-0.05 | -0.00284 | -0.00736 | 0.00169 | 0.51653 |
| | 0.5-0.05 | 0.00335 | -0.00118 | 0.00787 | 0.30292 |
| | 0.75-0.05 | 0.00798 | 0.00345 | 0.01250 | 0.00001 |
| | 1-0.05 | 0.02304 | 0.01851 | 0.02756 | 0.00000 |
| | 0.15-0.1 | -0.00227 | -0.00679 | 0.00225 | 0.76689 |
| | 0.25-0.1 | 0.00287 | -0.00165 | 0.00739 | 0.50189 |
| | 0.5-0.1 | 0.00905 | 0.00453 | 0.01357 | 0.00000 |
| | 0.75-0.1 | 0.01368 | 0.00916 | 0.01820 | 0.00000 |
| | 1-0.1 | 0.02874 | 0.02422 | 0.03326 | 0.00000 |
| | 0.25-0.15 | 0.00514 | 0.00062 | 0.00966 | 0.01496 |
| | 0.5-0.15 | 0.01132 | 0.00680 | 0.01585 | 0.00000 |
| | 0.75-0.15 | 0.01595 | 0.01143 | 0.02048 | 0.00000 |
| | 1-0.15 | 0.03101 | 0.02649 | 0.03554 | 0.00000 |
| | 0.5-0.25 | 0.00618 | 0.00166 | 0.01070 | 0.00146 |
| | 0.75-0.25 | 0.01081 | 0.00629 | 0.01534 | 0.00000 |
| | 1-0.25 | 0.02587 | 0.02135 | 0.03039 | 0.00000 |
| | 0.75-0.5 | 0.00463 | 0.00011 | 0.00915 | 0.04097 |
| | 1-0.5 | 0.01969 | 0.01517 | 0.02421 | 0.00000 |
| | 1-0.75 | 0.01506 | 0.01054 | 0.01958 | 0.00000 |

| | | | | | |
|---|---|---|---|---|---|
| **Extinction** | 0.05-0 | -0.06433 | -0.07768 | -0.05097 | 0.00000 |
| | 0.1-0 | -0.09113 | -0.10448 | -0.07777 | 0.00000 |
| | 0.15-0 | -0.10140 | -0.11476 | -0.08805 | 0.00000 |
| | 0.25-0 | -0.10701 | -0.12037 | -0.09365 | 0.00000 |
| | 0.5-0 | -0.11071 | -0.12407 | -0.09736 | 0.00000 |
| | 0.75-0 | -0.12143 | -0.13479 | -0.10808 | 0.00000 |
| | 1-0 | -0.13552 | -0.14887 | -0.12216 | 0.00000 |
| | 0.1-0.05 | -0.02680 | -0.04016 | -0.01344 | 0.00000 |
| | 0.15-0.05 | -0.03708 | -0.05043 | -0.02372 | 0.00000 |
| | 0.25-0.05 | -0.04268 | -0.05604 | -0.02933 | 0.00000 |
| | 0.5-0.05 | -0.04639 | -0.05974 | -0.03303 | 0.00000 |
| | 0.75-0.05 | -0.05711 | -0.07046 | -0.04375 | 0.00000 |
| | 1-0.05 | -0.07119 | -0.08455 | -0.05783 | 0.00000 |
| | 0.15-0.1 | -0.01028 | -0.02363 | 0.00308 | 0.25608 |
| | 0.25-0.1 | -0.01588 | -0.02924 | -0.00253 | 0.00909 |
| | 0.5-0.1 | -0.01959 | -0.03294 | -0.00623 | 0.00049 |
| | 0.75-0.1 | -0.03031 | -0.04366 | -0.01695 | 0.00000 |
| | 1-0.1 | -0.04439 | -0.05774 | -0.03103 | 0.00000 |
| | 0.25-0.15 | -0.00561 | -0.01896 | 0.00775 | 0.89211 |
| | 0.5-0.15 | -0.00931 | -0.02266 | 0.00405 | 0.37857 |
| | 0.75-0.15 | -0.02003 | -0.03339 | -0.00667 | 0.00034 |
| | 1-0.15 | -0.03411 | -0.04747 | -0.02075 | 0.00000 |
| | 0.5-0.25 | -0.00370 | -0.01706 | 0.00965 | 0.98819 |
| | 0.75-0.25 | -0.01442 | -0.02778 | -0.00107 | 0.02517 |
| | 1-0.25 | -0.02851 | -0.04186 | -0.01515 | 0.00000 |
| | 0.75-0.5 | -0.01072 | -0.02408 | 0.00263 | 0.20963 |
| | 1-0.5 | -0.02480 | -0.03816 | -0.01145 | 0.00000 |
| | 1-0.75 | -0.01408 | -0.02744 | -0.00072 | 0.03158 |

| | | | | | |
|---|---|---|---|---|---|
| **Diversification** | 0.05-0 | 0.08170 | 0.06822 | 0.09518 | 0.00000 |
| | 0.1-0 | 0.10280 | 0.08932 | 0.11628 | 0.00000 |
| | 0.15-0 | 0.11080 | 0.09732 | 0.12428 | 0.00000 |
| | 0.25-0 | 0.12155 | 0.10807 | 0.13503 | 0.00000 |
| | 0.5-0 | 0.13144 | 0.11796 | 0.14492 | 0.00000 |
| | 0.75-0 | 0.14679 | 0.13331 | 0.16027 | 0.00000 |
| | 1-0 | 0.17593 | 0.16245 | 0.18941 | 0.00000 |
| | 0.1-0.05 | 0.02109 | 0.00762 | 0.03457 | 0.00016 |
| | 0.15-0.05 | 0.02910 | 0.01562 | 0.04258 | 0.00000 |
| | 0.25-0.05 | 0.03985 | 0.02637 | 0.05333 | 0.00000 |
| | 0.5-0.05 | 0.04973 | 0.03625 | 0.06321 | 0.00000 |
| | 0.75-0.05 | 0.06508 | 0.05161 | 0.07856 | 0.00000 |
| | 1-0.05 | 0.09422 | 0.08075 | 0.10770 | 0.00000 |
| | 0.15-0.1 | 0.00801 | -0.00547 | 0.02148 | 0.58614 |
| | 0.25-0.1 | 0.01875 | 0.00527 | 0.03223 | 0.00113 |
| | 0.5-0.1 | 0.02864 | 0.01516 | 0.04212 | 0.00000 |
| | 0.75-0.1 | 0.04399 | 0.03051 | 0.05747 | 0.00000 |
| | 1-0.1 | 0.07313 | 0.05965 | 0.08661 | 0.00000 |
| | 0.25-0.15 | 0.01075 | -0.00273 | 0.02423 | 0.21690 |
| | 0.5-0.15 | 0.02063 | 0.00715 | 0.03411 | 0.00023 |
| | 0.75-0.15 | 0.03598 | 0.02251 | 0.04946 | 0.00000 |
| | 1-0.15 | 0.06512 | 0.05164 | 0.07860 | 0.00000 |
| | 0.5-0.25 | 0.00989 | -0.00359 | 0.02336 | 0.31370 |
| | 0.75-0.25 | 0.02524 | 0.01176 | 0.03872 | 0.00000 |
| | 1-0.25 | 0.05438 | 0.04090 | 0.06786 | 0.00000 |
| | 0.75-0.5 | 0.01535 | 0.00187 | 0.02883 | 0.01469 |
| | 1-0.5 | 0.04449 | 0.03101 | 0.05797 | 0.00000 |
| | 1-0.75 | 0.02914 | 0.01566 | 0.04262 | 0.00012 |

**Table 4: Tukey's test results correspond to simulations that verify the effect of phenotypic plasticity on adaptive rates in the context of fast climatic changes and presence of the plasticity cost**

| | | | 95% Confidence Interval | | |
|---|---|---|---|---|---|
| **Variables** | Factor levels | Mean difference | Lower bound | Upper bound | Significance |

| | | | | | |
|---|---|---|---|---|---|
| **Trait evolution** | 0.05-0 | -0.00027 | -0.00037 | -0.00017 | 0.00000 |
| | 0.1-0 | -0.00034 | -0.00044 | -0.00024 | 0.00000 |
| | 0.15-0 | -0.00042 | -0.00052 | -0.00031 | 0.00000 |
| | 0.25-0 | -0.00043 | -0.00054 | -0.00033 | 0.00000 |
| | 0.5-0 | -0.00042 | -0.00052 | -0.00032 | 0.00000 |
| | 0.75-0 | -0.00047 | -0.00057 | -0.00037 | 0.00000 |
| | 1-0 | -0.00054 | -0.00064 | -0.00044 | 0.00000 |
| | 0.1-0.05 | -0.00006 | -0.00017 | 0.00004 | 0.49449 |
| | 0.15-0.05 | -0.00014 | -0.00024 | -0.00004 | 0.00097 |
| | 0.25-0.05 | -0.00016 | -0.00026 | -0.00006 | 0.00012 |
| | 0.5-0.05 | -0.00015 | -0.00025 | -0.00005 | 0.00043 |
| | 0.75-0.05 | -0.00019 | -0.00030 | -0.00009 | 0.00000 |
| | 1-0.05 | -0.00027 | -0.00037 | -0.00017 | 0.00000 |
| | 0.15-0.1 | -0.00008 | -0.00018 | 0.00002 | 0.25978 |
| | 0.25-0.1 | -0.00010 | -0.00020 | 0.00000 | 0.07302 |
| | 0.5-0.1 | -0.00008 | -0.00019 | 0.00002 | 0.16650 |
| | 0.75-0.1 | -0.00013 | -0.00023 | -0.00003 | 0.00380 |
| | 1-0.1 | -0.00020 | -0.00030 | -0.00010 | 0.00000 |
| | 0.25-0.15 | -0.00002 | -0.00012 | 0.00008 | 0.99902 |
| | 0.5-0.15 | -0.00001 | -0.00011 | 0.00009 | 0.99999 |
| | 0.75-0.15 | -0.00005 | -0.00015 | 0.00005 | 0.75465 |
| | 1-0.15 | -0.00012 | -0.00023 | -0.00002 | 0.00636 |
| | 0.5-0.25 | 0.00001 | -0.00009 | 0.00011 | 0.99996 |
| | 0.75-0.25 | -0.00003 | -0.00013 | 0.00007 | 0.97143 |
| | 1-0.25 | -0.00011 | -0.00021 | -0.00000 | 0.03576 |
| | 0.75-0.5 | -0.00004 | -0.00015 | 0.00006 | 0.86917 |
| | 1-0.5 | -0.00012 | -0.00022 | -0.00002 | 0.01276 |
| | 1-0.75 | -0.00007 | -0.00017 | 0.00003 | 0.34341 |

| | | | | | |
|---|---|---|---|---|---|
| **Speciation** | 0.05-0<br>0.1-0<br>0.15-0<br>0.25-0<br>0.5-0<br>0.75-0<br>1-0<br>0.1-0.05<br>0.15-0.05<br>0.25-0.05<br>0.5-0.05<br>0.75-0.05<br>1-0.05<br>0.15-0.1<br>0.25-0.1<br>0.5-0.1<br>0.75-0.1<br>1-0.1<br>0.25-0.15<br>0.5-0.15<br>0.75-0.15<br>1-0.15<br>0.5-0.25<br>0.75-0.25<br>1-0.25<br>0.75-0.5<br>1-0.5<br>1-0.75 | 0.01696<br>0.01326<br>0.01342<br>0.01512<br>0.01493<br>0.02501<br>0.04041<br>-0.00370<br>-0.00354<br>-0.00184<br>-0.00203<br>0.00805<br>0.02345<br>0.00015<br>0.00186<br>0.00166<br>0.01175<br>0.02715<br>0.00171<br>0.00151<br>0.01160<br>0.02699<br>-0.00020<br>0.00989<br>0.02529<br>0.01008<br>0.02549<br>0.01540 | 0.01201<br>0.00831<br>0.00846<br>0.01017<br>0.00997<br>0.02006<br>0.03546<br>-0.00865<br>-0.00850<br>-0.00679<br>-0.00699<br>0.00310<br>0.01850<br>-0.00480<br>-0.00309<br>-0.00329<br>0.00679<br>0.02220<br>-0.00324<br>-0.00344<br>0.00664<br>0.02204<br>-0.00515<br>0.00494<br>0.02034<br>0.00513<br>0.02053<br>0.01044 | 0.02191<br>0.01822<br>0.01837<br>0.02008<br>0.01988<br>0.02997<br>0.04537<br>0.00126<br>0.00141<br>0.00312<br>0.00292<br>0.01301<br>0.02841<br>0.00511<br>0.00681<br>0.00662<br>0.01670<br>0.03210<br>0.00666<br>0.00646<br>0.01655<br>0.03195<br>0.00476<br>0.01484<br>0.03024<br>0.01504<br>0.03044<br>0.02035 | 0.00000<br>0.00000<br>0.00000<br>0.00000<br>0.00000<br>0.00000<br>0.00000<br>0.29237<br>0.34495<br>0.94092<br>0.90283<br>0.00008<br>0.00000<br>1.00000<br>0.93743<br>0.96504<br>0.00000<br>0.00000<br>0.96000<br>0.97953<br>0.00000<br>0.00000<br>1.00000<br>0.00001<br>0.00000<br>0.00000<br>0.00000<br>0.00000 |

| | | | | | |
|---|---|---|---|---|---|
| **Extinction** | 0.05-0 | -0.06314 | -0.07703 | -0.04924 | 0.00000 |
| | 0.1-0 | -0.08454 | -0.09844 | -0.07065 | 0.00000 |
| | 0.15-0 | -0.09528 | -0.10918 | -0.08139 | 0.00000 |
| | 0.25-0 | -0.10590 | -0.11980 | -0.09201 | 0.00000 |
| | 0.5-0 | -0.11190 | -0.12580 | -0.09801 | 0.00000 |
| | 0.75-0 | -0.11641 | -0.13031 | -0.10252 | 0.00000 |
| | 1-0 | -0.13552 | -0.14941 | -0.12162 | 0.00000 |
| | 0.1-0.05 | -0.02141 | -0.03530 | -0.00751 | 0.00021 |
| | 0.15-0.05 | -0.03214 | -0.04604 | -0.01825 | 0.00000 |
| | 0.25-0.05 | -0.04276 | -0.05666 | -0.02887 | 0.00000 |
| | 0.5-0.05 | -0.04877 | -0.06266 | -0.03487 | 0.00000 |
| | 0.75-0.05 | -0.05328 | -0.06717 | -0.03938 | 0.00000 |
| | 1-0.05 | -0.07238 | -0.08627 | -0.05848 | 0.00000 |
| | 0.15-0.1 | -0.01074 | -0.02463 | 0.00316 | 0.25136 |
| | 0.25-0.1 | -0.02136 | -0.03525 | -0.00746 | 0.00022 |
| | 0.5-0.1 | -0.02736 | -0.04125 | -0.01346 | 0.00000 |
| | 0.75-0.1 | -0.03187 | -0.04576 | -0.01797 | 0.00000 |
| | 1-0.1 | -0.05097 | -0.06487 | -0.03708 | 0.00000 |
| | 0.25-0.15 | -0.01062 | -0.02451 | 0.00327 | 0.26395 |
| | 0.5-0.15 | -0.01662 | -0.03052 | -0.00273 | 0.00849 |
| | 0.75-0.15 | -0.02113 | -0.03503 | -0.00724 | 0.00026 |
| | 1-0.15 | -0.04023 | -0.05413 | -0.02634 | 0.00000 |
| | 0.5-0.25 | -0.00600 | -0.01990 | 0.00789 | 0.87674 |
| | 0.75-0.25 | -0.01051 | -0.02441 | 0.00338 | 0.27607 |
| | 1-0.25 | -0.02961 | -0.04351 | -0.01572 | 0.00000 |
| | 0.75-0.5 | -0.00451 | -0.01840 | 0.00939 | 0.97101 |
| | 1-0.5 | -0.02361 | -0.03751 | -0.00972 | 0.00003 |
| | 1-0.75 | -0.01910 | -0.03299 | -0.00521 | 0.00134 |

| | | | | | |
|---|---|---|---|---|---|
| **Diversification** | 0.05-0 | 0.08010 | 0.06649 | 0.09370 | 0.00000 |
| | 0.1-0 | 0.09781 | 0.08420 | 0.11141 | 0.00000 |
| | 0.15-0 | 0.10870 | 0.09509 | 0.12230 | 0.00000 |
| | 0.25-0 | 0.12102 | 0.10742 | 0.13463 | 0.00000 |
| | 0.5-0 | 0.12683 | 0.11322 | 0.14044 | 0.00000 |
| | 0.75-0 | 0.14143 | 0.12782 | 0.15503 | 0.00000 |
| | 1-0 | 0.17593 | 0.16232 | 0.18953 | 0.00000 |
| | 0.1-0.05 | 0.01771 | 0.00410 | 0.03132 | 0.00293 |
| | 0.15-0.05 | 0.02860 | 0.01499 | 0.04221 | 0.00000 |
| | 0.25-0.05 | 0.04093 | 0.02731 | 0.05453 | 0.00000 |
| | 0.5-0.05 | 0.04673 | 0.03313 | 0.06034 | 0.00000 |
| | 0.75-0.05 | 0.06133 | 0.04772 | 0.07494 | 0.00000 |
| | 1-0.05 | 0.09583 | 0.08223 | 0.10944 | 0.00000 |
| | 0.15-0.1 | 0.01089 | -0.00272 | 0.02450 | 0.21279 |
| | 0.25-0.1 | 0.02322 | 0.00961 | 0.03682 | 0.00003 |
| | 0.5-0.1 | 0.02902 | 0.01542 | 0.04263 | 0.00000 |
| | 0.75-0.1 | 0.04362 | 0.03001 | 0.05723 | 0.00000 |
| | 1-0.1 | 0.07812 | 0.06452 | 0.09173 | 0.00000 |
| | 0.25-0.15 | 0.01233 | -0.00128 | 0.02593 | 0.10409 |
| | 0.5-0.15 | 0.01813 | 0.00453 | 0.03174 | 0.00211 |
| | 0.75-0.15 | 0.03273 | 0.01912 | 0.04634 | 0.00000 |
| | 1-0.15 | 0.06723 | 0.05363 | 0.08084 | 0.00000 |
| | 0.5-0.25 | 0.00581 | -0.00780 | 0.01941 | 0.88335 |
| | 0.75-0.25 | 0.02040 | 0.00680 | 0.03401 | 0.00034 |
| | 1-0.25 | 0.05491 | 0.04130 | 0.06851 | 0.00000 |
| | 0.75-0.5 | 0.01460 | 0.00099 | 0.02820 | 0.02683 |
| | 1-0.5 | 0.04910 | 0.03549 | 0.06271 | 0.00000 |
| | 1-0.75 | 0.03450 | 0.02090 | 0.04811 | 0.00000 |

51
